# GluN2B-containing NMDA receptors are required for potentiation and depression of responses in ocular dominance plasticity

**DOI:** 10.1101/2022.12.01.518760

**Authors:** Michelle Bridi, Su Hong, Daniel Severin, Alfredo Kirkwood

**Affiliations:** Zanvyl Krieger Mind/Brain Institute, Johns Hopkins University, Baltimore, MD 21218; Rockefeller Neuroscience Institute, West Virginia University, Morgantown, WV 26506

## Abstract

Monocular deprivation (MD) causes an initial decrease in synaptic responses to the deprived eye in juvenile mouse primary visual cortex (V1) through Hebbian long-term depression (LTD). This is followed by a homeostatic increase, which has been attributed to synaptic scaling. However, homeostasis during other forms of visual deprivation is caused by sliding the threshold for Hebbian long-term potentiation (LTP) rather than scaling. We therefore asked whether the homeostatic increase during MD requires GluN2B-containing NMDA receptor activity, which is required to slide the plasticity threshold but not for synaptic scaling. Selective GluN2B blockade from 2-6d after monocular lid suture prevented the homeostatic increase in miniature excitatory postsynaptic current (mEPSC) amplitude in monocular V1 of acute slices and prevented the increase in visually evoked responses in binocular V1 *in vivo*. The decrease in mEPSC amplitude and visually evoked responses during the first 2d of MD also required GluN2B activity. Together, these results indicate that GluN2B-containing NMDA receptors first play a role in LTD immediately following eye closure, and then promote homeostasis during prolonged MD by sliding the plasticity threshold in favor of LTP.

## Introduction

Maintaining neuronal firing rates within an optimal range is considered essential for proper neuronal processing, and can be achieved by changing postsynaptic strength to compensate for altered spiking activity (e.g. as a result of sensory deprivation) (reviewed in Davis, 2006). For example, visual deprivation by dark exposure, intraocular tetrodotoxin injection, eyelid suture, enucleation, and retinal lesions increase synaptic strength in primary visual cortex (V1) (Desai et al., 2002; Gao et al., 2010; Goel and Lee, 2007; Goel et al., 2011; He et al., 2012; Hengen et al., 2013; Keck et al., 2013; Lambo and Turrigiano, 2013). Similarly, deafening and whisker trimming increase synaptic strength in primary auditory and somatosensory cortices, respectively (Glazewski et al., 2017; Kotak et al., 2005). However, the exact mechanisms by which homeostatic synaptic strengthening occur remain unclear.

Two distinct mechanisms that can homeostatically increase postsynaptic strength are the sliding threshold for plasticity and synaptic scaling. Lowering the plasticity threshold enables synaptic strengthening via Hebbian long-term potentiation (LTP), whereas non-Hebbian synaptic scaling globally increases the strength of all synapses by the same factor (Abraham and Bear, 1996; Turrigiano et al., 1998). These mechanisms can be distinguished based on their dependence on GluN2B-containing NMDA receptors: increased GluN2B levels lower the threshold for Hebbian plasticity, but are not required for synaptic scaling (Philpot et al., 2001; Turrigiano et al., 1998). Previous studies indicate that visual deprivation by dark exposure increases synaptic strength in V1 by lowering the threshold for Hebbian LTP (Bridi et al., 2018; Rodriguez et al., 2019), but whether this is true for other forms of visual deprivation is unknown.

Monocular lid suture (monocular deprivation, MD) shifts the visual responsiveness of V1 in favor of the open eye, in a process referred to as ocular dominance plasticity (ODP). In juvenile mice, MD causes an initial (within 3d) decrease in visually evoked cortical responses to the deprived eye, followed by a delayed (5-7d) homeostatic increase in responses to both eyes (Frenkel and Bear, 2004; Kaneko et al., 2008; Ranson et al., 2012). These changes are paralleled by an initial decrease and delayed increase in postsynaptic strength in V1 layer 2/3 pyramidal neurons, which have been interpreted as Hebbian long-term depression (LTD) and synaptic scaling, respectively (Lambo and Turrigiano, 2013). However, GluN2B-containing NMDA receptor levels increase during the delayed phase of ODP (Chen and Bear, 2007; Cooper and Bear, 2012; Guo et al., 2017), suggesting that prolonged MD may increase synaptic strength by decreasing the threshold for LTP.

We therefore wished to test whether homeostasis during late ODP was caused by the sliding threshold rather than synaptic scaling. To this end, we administered a GluN2B-specific antagonist specifically during the late phase of ODP. This manipulation prevented the homeostatic increases in both mEPSC amplitude and visually evoked responses in V1. We also investigated the mechanisms of plasticity during early ODP and found that GluN2B-containing NMDA receptors are required for the initial weakening of synapses and evoked responses in V1. Together, our findings indicate that MD initially weakens cortical synapses via GluN2B-dependent LTD, then homeostatically increases synaptic strength by lowering the threshold for Hebbian LTP.

## Methods

### Animals

C57BL/6J mice were raised (no more than 5 per cage) on a 12:12 light:dark cycle with food and water *ad libitum*. Equal numbers of male and female mice were used in each group. All procedures conform to the guidelines of the U.S. Department of Health and Human Services Office of Laboratory Animal Welfare (OLAW) and were approved by the Johns Hopkins University Institutional Animal Care and Use Committee. Sample sizes were chosen to correspond with previous studies in which the effects of visual manipulation were measured. For each experiment, animals within a litter were randomly distributed across groups.

### Monocular deprivation

Naïve mice underwent monocular lid suture beginning at postnatal day 25-27, and mice were age-matched across groups in all experiments. Monocular lid suture was performed under isoflurane anesthesia. For brief (2d) MD, the right eyelids were sutured together using 7-0 polypropylene suture no more than 48h prior to the experiment. For 6d MD, the margins of the upper and lower lids were trimmed prior to suture. Neosporin was applied to prevent infection. Animals were disqualified in the event of suture opening or infection.

### Drug administration

Osmotic minipumps (Alzet 1007D, Durect Corp., Cupertino, CA) were filled with vehicle (20%DMSO/80% saline) or Ro 25-6981 (Cayman Chemical, Ann Arbor, MI) at a concentration to deliver 30 mg/kg/day. The pump was primed in 0.9% NaCl at 37°C for at least 6h prior to implantation. The minipump was implanted subcutaneously under isoflurane anesthesia; the incision was sutured shut and Neosporin was applied to prevent infection. Meloxicam (5mg/kg, s.c.) was administered to reduce pain and inflammation.

### Slice electrophysiology

Visual cortical slices were prepared as previously described (Guo et al., 2012). 300 μm thick slices were cut in ice-cold dissection buffer containing (in mM): 212.7 sucrose, 5 KCl, 1.25 NaH_2_PO_4_, 10 MgCl_2_, 0.5 CaCl_2_, 26 NaHCO_3_, 10 dextrose, bubbled with 95% O2/5% CO_2_ (pH 7.4). Slices were transferred to artificial cerebrospinal fluid (ACSF) containing (in mM): 124 NaCl, 5 KCl, 1.25 NaH_2_PO_4_, 1 MgCl_2_, 2 CaCl_2_, 26 NaHCO_3_, 10 dextrose, bubbled with 95% O2/5% CO_2_ (pH 7.4). Slices were incubated in ACSF at 30°C for 30 minutes, then at room temperature for at least 30 minutes prior to recording. Visualized whole-cell recordings were obtained from pyramidal neurons in V1 layer 2/3 using 3-6 MΩ glass pipettes. Data were filtered at 2 kHz and digitized at 10 kHz using Igor Pro (WaveMetrics Inc., Lake Oswego, OR). Cells were excluded if input or series resistance changed >20% during the recording.

### Miniature EPSC recordings

Miniature EPSCs were recorded from pyramidal cells in layer 2/3 of V1m. Cells were recorded in both hemispheres, with non-deprived hemisphere (ipsilateral to the deprived eye) serving as control. Recordings were performed with an intracellular solution containing (in mM): 130 Cs-gluconate, 8 KCl, 1 EGTA, 10 HEPES, 4 (Na)ATP, 5 QX-314 (pH adjusted to 7.25 with CsOH, 280-290 mOsm) under voltage clamp (Vh: -70 mV). 1 μM TTX, 100μM APV, and 2.5 μM gabazine were included in the bath. Events were detected and analyzed using Mini Analysis (Synaptosoft, Decatur, GA). Cells with root mean square (RMS) of membrane current noise < 2, input resistance >200MΩ, and series resistance <20 MΩ were included in the analysis. The threshold for mini detection was set at three times the RMS noise. The first 300 non-overlapping events with rise times ≤ 3 ms were used to estimate the mEPSC amplitude distribution and produce the average mEPSC for the cell. Cells were examined for dendritic filtering by confirming that there was not a negative correlation between mEPSC amplitude and rise time.

### NMDA receptor current recordings

NMDA receptor currents were recorded from V1m of both hemispheres, with the right hemisphere serving as control, or in the binocular zone of the left (deprived) hemisphere, with normally reared animals serving as control. Recordings were made using an internal pipette solution containing (in mM): 102 Cs-gluconate, 5 TEA-chloride, 3.7 NaCl, 20 HEPES, 0.3 (Na)GTP, 4 (Mg)ATP, 0.2 EGTA, 10 BAPTA, 5 QX-314 (pH 7.2, ∼300 mOsm) under voltage clamp (V_h_=+40 mV). To isolate NMDA receptor currents and minimize multisynaptic responses, ACSF in the recording chamber contained 2.5 μM gabazine, 25μM CNQX, 1μM glycine, 4mM CaCl_2_, and 4mM MgCl_2_. A concentric bipolar electrode (FHC, Bowdoin, ME) was placed in the middle of the cortical thickness. Stimulation intensity was set to evoke responses of at least 100 pA, and slices were stimulated every 15 seconds. Traces free of noise were averaged and NMDA receptor deactivation kinetics were measured by fitting a double exponential function to the current decay of the averaged trace (Igor Pro). The weighted decay constant, 1_w_, was calculated as: 1_w_=1_f_(I_f_/(I_f_+I_s_))+1_s_(I_s_/(I_f_+I_s_))(Guo et al., 2012; Rumbaugh and Vicini, 1999).

### Optical Imaging of the Intrinsic Cortical Signal

Isoflurane in O_2_ (2-3% for induction, 0.5-1% for maintenance) was supplemented with chlorprothixene (2 mg/kg i.p.). Hair was removed using depilatory cream and the scalp was sterilized with povidone iodine. An incision was made in the scalp and lidocaine was applied to the incision margins. The exposed skull above V1 contralateral to the deprived eye was covered in 3% agarose and an 8mm round glass coverslip. The surface vasculature and intrinsic signals were imaged using a Dalsa 1M30 CCD camera (Dalsa, Waterloo, Canada). Surface vasculature was visualized by illuminating the area with 555nm light. Then the camera was focused 600 μm below the cortical surface and the area was illuminated with 610 nm light. A high refresh rate monitor (1024 × 768 @120 Hz; ViewSonic, Brea, CA) was aligned in the center of the mouse’s visual field 25cm in front of the eyes. Visual stimuli consisted of a white horizontal bar (2° height, 20° width) restricted to the binocular visual field (-5° to +15° azimuth), on a black background. Each eye was individually presented with the stimulus moving continuously in the upward and downward direction for 7 minutes per direction. The cortical response at the stimulus frequency was extracted by Fourier analysis and maps generated for each eye were averaged. The maps for each eye were then analyzed (Matlab, Mathworks, Natick, MA). Images were smoothed by a 5×5 low-pass Gaussian filter and the binocular region of interest (ROI) was defined as the 70% of pixels with the highest intensity in the ipsilateral eye map. The average intensity at all pixels in the ROI was calculated for the ipsilateral and contralateral maps. For each pixel an ocular dominance value was calculated as (contra-ipsi)/(contra+ipsi) and all ocular dominance values in the ROI were averaged to obtain the ODI.

### Statistical Analysis

Normality was determined using the d’Agostino test. Groups with normally distributed data were compared using 2-tailed paired or unpaired *t* tests, one-way ANOVAs, or one-way repeated measures ANOVAs, as indicated. Holm-Sidak post-hoc tests were used for multiple comparisons following one-way ANOVAs. Groups that were not normally distributed were compared using nonparametric Mann-Whitney tests or ANOVAs on ranks followed by Dunn’s post-hoc test for multiple comparisons (GraphPad Prism, San Diego, CA). Statistical outliers were detected (ROUT test) using pre-established criteria and excluded from analysis.

## Results

### GluN2B is required for homeostasis in V1m

In monocular V1 (V1m) contralateral to the deprived eye, 2d MD decreases mEPSC amplitude, followed by a homeostatic increase by 6d (Lambo and Turrigiano, 2013). 6d MD also increases the GluN2B component of the NMDA receptor response (Guo et al., 2017). We wished to determine whether this increase in GluN2B-containing NMDA receptors is required for the increase in mEPSC amplitude. To this end, we recorded mEPSCs in V1m of acute slices obtained from normal reared (NR) mice and mice that had undergone MD (Fig. 1A). In agreement with previous findings (Lambo and Turrigiano, 2013), 2 days of MD decreased mEPSC amplitude in the deprived (contralateral) hemisphere (Fig. 1B, C; Table 1). We then tested whether increased mEPSC amplitude during late-phase MD (between 2 and 6 days) required activation of GluN2B-containing NMDA receptors. After 2d MD, an osmotic minipump containing the GluN2B-specific antagonist Ro 25-6981 or its vehicle was implanted subcutaneously for the remainder of the MD period (Fig. 1A). mEPSC amplitude increased above control levels in animals that received vehicle infusions (Fig. 1B, C). However, this increase was prevented by blocking GluN2B-contatining NMDA receptors (Fig. 1B, C; Table 1). This GluN2B dependence is consistent with the sliding threshold mechanism underlying the homeostatic increase in mEPSC amplitude during MD.

**Table 1.** mEPSC properties and recording conditions in NR, 2d MD, 6dMD+vehicle, and 6dMD+Ro 25-6981 groups (related to Figure 1). For each treatment, data were compared between control (ipsi) and deprived (contra) hemispheres using 2-tailed unpaired *t* tests (*t* statistic reported) or Mann-Whitney tests (*U* statistic reported), as indicated. Bold values indicate significant differences. Data are shown as mean±SEM. N is reported as cells, mice.

|  |  | Amplitude<br>(pA) | Frequency<br>(Hz) | Rise<br>(ms) | Decay<br>(ms) | Rin<br>(MW) | Rs<br>(MW) | RMS | N |
| --- | --- | --- | --- | --- | --- | --- | --- | --- | --- |
| NR | Ipsi | 13.3 ± 0.4 | 7.87 ± 0.6 | 0.76 ± 0.02 | 2.56 ± 0.1 | 273 ± 13 | 16.8 ± 0.3 | 1.84 ± 0.02 | 27, 5 |
|  | Contra | 13.3 ± 0.4 | 8.90 ± 0.6 | 0.75 ± 0.02 | 2.56 ± 0.1 | 294 ± 12 | 16.2 ± 0.3 | 1.82 ± 0.02 | 29, 5 |
| | statistic | $t_{(54)}=0.06$ | $t_{(54)}=1.2$ | $U=390.5$ | $t_{(54)}=0.03$ | $U=289$ | $t_{(54)}=1.2$ | $U=358.5$ | |
|  | *P* | 0.95 | 0.22 | 0.99 | 0.97 | 0.09 | 0.2 | 0.59 |  |
| 2d MD | Ipsi | 13.4 ± 0.3 | 7.00 ± 0.5 | 0.81 ± 0.02 | 2.72 ± 0.1 | 283 ± 15 | 16.5 ± 0.4 | 1.83 ± 0.02 | 25, 5 |
|  | Contra | 12.3 ± 0.3 | 9.76 ± 0.7 | 0.79 ± 0.02 | 2.69 ± 0.1 | 278 ± 17 | 16.6 ± 0.4 | 1.81 ± 0.02 | 27, 5 |
| | statistic | $t_{(50)}=2.5$ | $t_{(50)}=3.1$ | $t_{(50)}=0.83$ | $t_{(50)}=0.37$ | $U=313.5$ | $t_{(50)}=0.2$ | $t_{(50)}=0.63$ | |
|  | *P* | **0.018\*** | **0.003\*** | 0.41 | 0.71 | 0.67 | 0.8 | 0.53 |  |
| 6d MD<br>Vehicle | Ipsi | 12.3 ± 0.4 | 6.69 ± 0.5 | 0.81 ± 0.02 | 2.67 ± 0.1 | 307 ± 33 | 16.0 ± 0.4 | 1.81 ± 0.02 | 23, 5 |
|  | Contra | 13.4 ± 0.3 | 6.98 ± 0.5 | 0.78 ± 0.02 | 2.70 ± 0.1 | 271 ± 14 | 16.6 ± 0.3 | 1.81 ± 0.02 | 28, 5 |
| | statistic | $t_{(49)}=2.6$ | $U=308$ | $t_{(49)}=0.81$ | $U=305$ | $U=258.5$ | $t_{(49)}=1.2$ | $t_{(49)}=0.007$ | |
|  | *P* | **0.012\*** | 0.80 | 0.42 | 0.75 | 0.23 | 0.26 | 0.99 |  |
| 6d MD<br>Ro 25-6981 | Ipsi | 13.8 ± 0.3 | 7.40 ± 0.5 | 0.82 ± 0.03 | 2.77 ± 0.1 | 282 ± 13 | 16.1 ± 0.4 | 1.80 ± 0.02 | 24, 5 |
|  | Contra | 12.4 ± 0.4 | 7.49 ± 0.6 | 0.85 ± 0.02 | 2.89 ± 0.1 | 318 ± 19 | 16.9 ± 0.5 | 1.81 ± 0.02 | 27, 5 |
| | statistic | $t_{(49)}=2.8$ | $t_{(49)}=0.11$ | $t_{(49)}=0.92$ | $t_{(49)}=1.09$ | $U=249.5$ | $t_{(49)}=1.1$ | $t_{(49)}=0.30$ | |
|  | *P* | **0.0075\*** | 0.91 | 0.36 | 0.28 | 0.16 | 0.3 | 0.77 |  |

**Figure 1.**
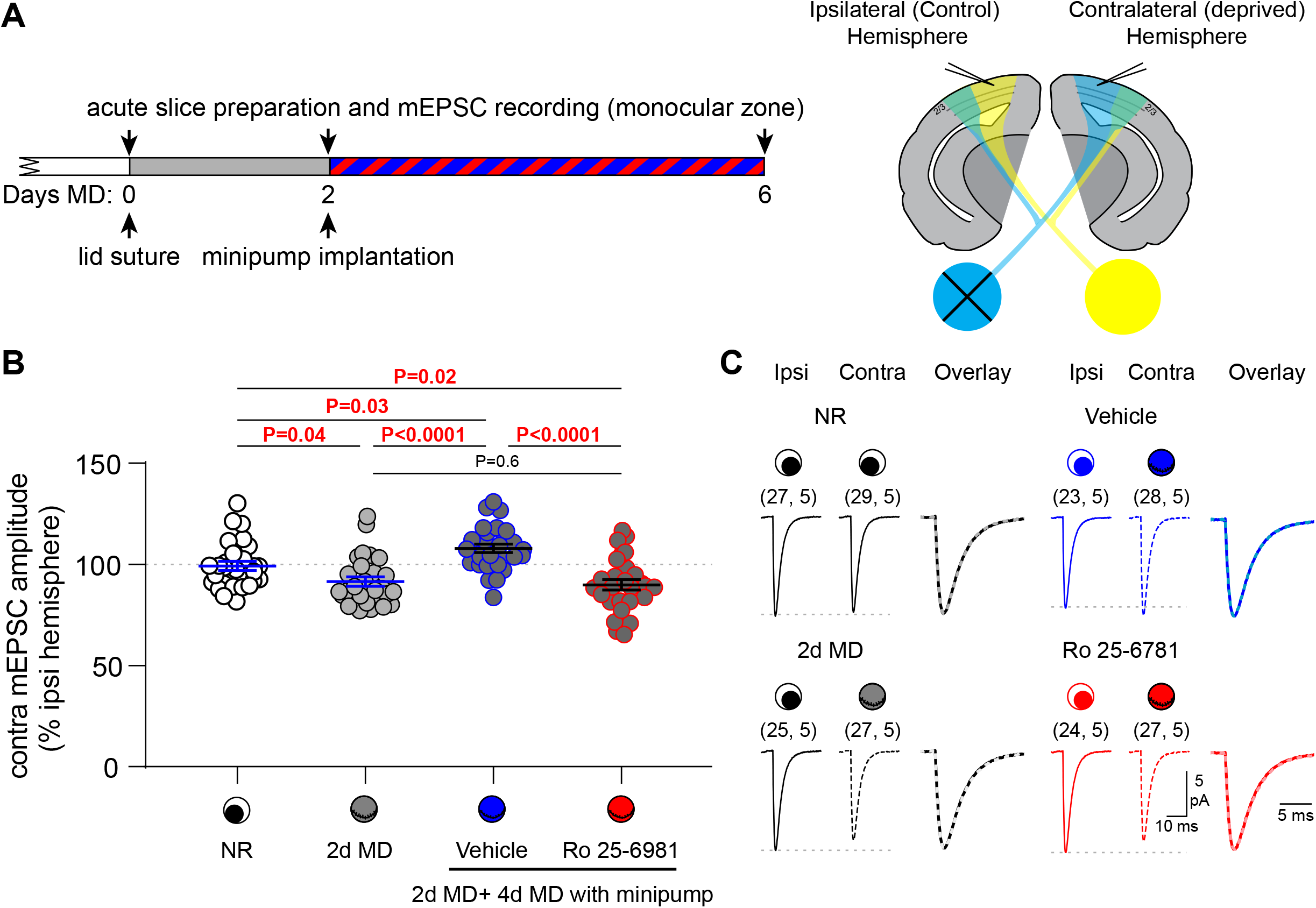
GluN2B-containing NMDA receptors are required for homeostasis during late-phase ocular dominance plasticity. (A) Experimental design. The right eyelid was sutured shut to initiate an ocular dominance shift. After two days, a minipump containing the GluN2B-specific antagonist Ro 25-6981 or its vehicle was implanted subcutaneously. Arrowheads indicate times at which acute brain slices were collected for mEPSC recording in the monocular zone of each hemisphere. (B) mEPSC amplitude in the contralateral (deprived) hemisphere is expressed as percentage of ipsilateral (control) hemisphere in the same animal. Left: compared to normally reared (NR) animals, mEPSC amplitude decreased after 2d MD and then increased above baseline levels after 6d MD with vehicle. Ro 25-6981 prevented mEPSC amplitude from increasing (ANOVA *F*_(3, 106)_=12.8, *P*<0.0001; Holm-Sidak post-hoc *P* values indicated). Data are shown as mean±SEM and sample size is indicated as (cells, mice). (C) Averaged mEPSC traces from each hemisphere. Traces normalized to peak amplitude (overlay) show no difference in kinetics (see Table 1).

### GluN2B is required for homeostasis in V1b

In binocular V1 (V1b), the amplitude of visually evoked deprived eye responses decreases during early MD, followed by an increase of responses to both eyes during late MD *in vivo* (Frenkel and Bear, 2004; Kaneko et al., 2008; Ranson et al., 2012). We wished to test whether the sliding threshold underlies the late increase in visual responses in V1b. We performed optical imaging of the intrinsic cortical signal in the same mice at multiple time points: baseline, 2d MD, and 6d MD (Fig. 2A). After the 2d MD imaging session, we implanted an osmotic minipump containing Ro 25-6981 or its vehicle. We measured the magnitude of cortical response to visual stimulation of each eye, and calculated the ocular dominance index at each time point (Fig. 2B-D). 2d MD induced an initial shift in ocular dominance that was caused by decreased response to the contralateral (deprived) eye, with no change in the response of the ipsilateral eye. During late-phase MD (between 2d and 6d), the magnitude of the cortical response to both the ipsilateral and contralateral eyes increased in the vehicle group, whereas Ro 25-6981 prevented this increase. We confirmed that chronic administration of Ro 25-6981 alone did not weaken responses in nondeprived V1 (Fig. 3). Together, these results indicate that the sliding threshold is required for homeostasis during late-phase MD.

**Figure 2.**
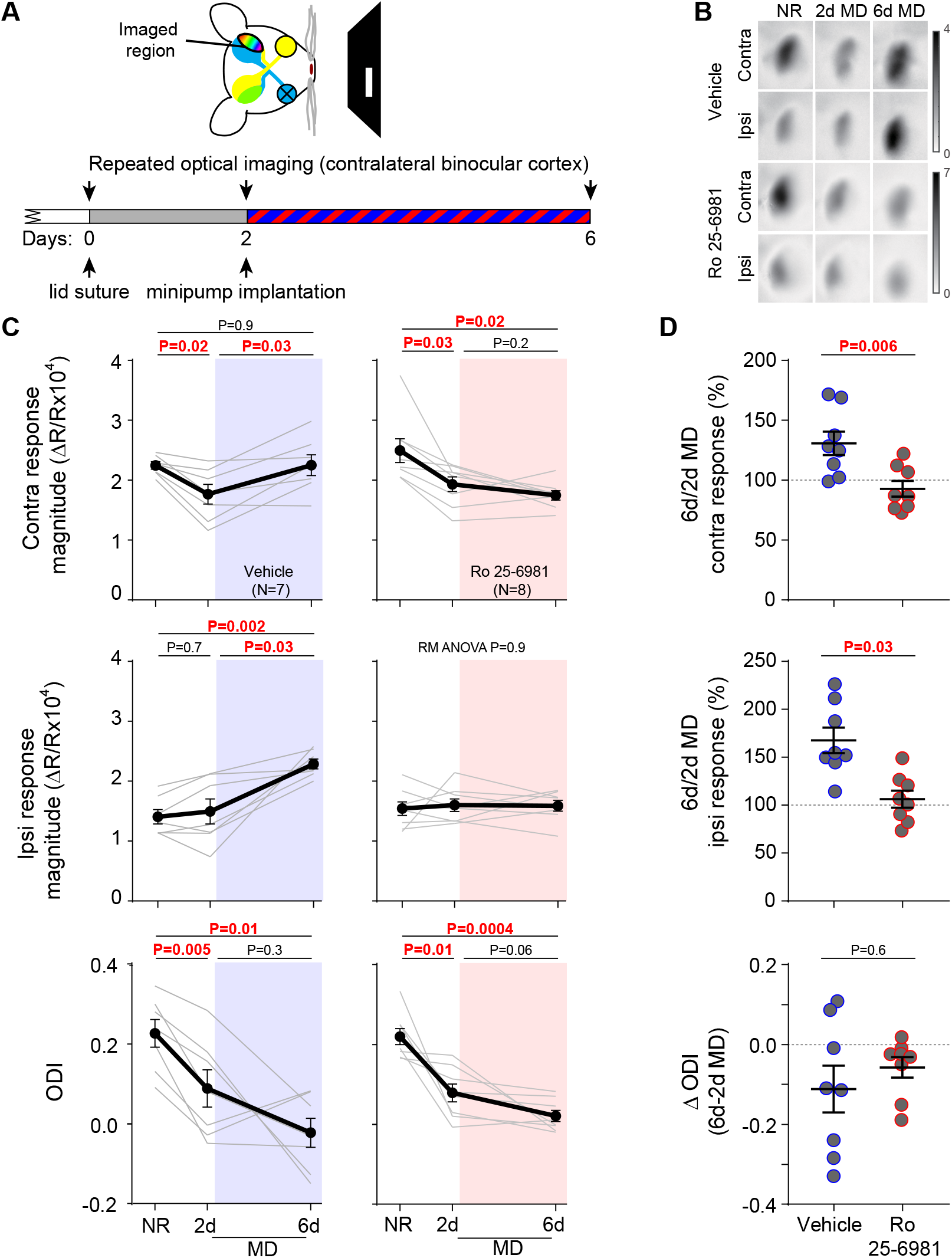
GluN2B-containing NMDA receptors are required for homeostasis during late-phase ocular dominance plasticity. (A) Experimental design. The right eyelid was sutured shut to initiate an ocular dominance shift. After two days, a minipump containing the GluN2B-specific antagonist Ro 25-6981 or its vehicle was implanted subcutaneously. Arrowheads indicate optical imaging sessions. The same animals were imaged at all three time points. (B) Representative images. (C) In both groups, there was an initial decrease in ODI at 2d MD (Vehicle: RM ANOVA *F*_(1.2,7.4)_=10.5, *P*=0.01; Ro: RM ANOVA *F*_(1.6, 10.9)_=22.4, *P*=0.0002). This was caused by decreased contralateral eye responses (Vehicle: RM ANOVA *F*_(1.8, 10.7)_=7.6, *P*=0.01; Ro: RM ANOVA *F*_(1.7,11.8)_=11, *P*=0.003). Ro 25-6981 treatment over the subsequent 4d MD blocked the increase in contralateral and ipsilateral response magnitude (Vehicle ipsi: RM ANOVA *F*_(1.2, 7)_=15.7, *P*=0.005; Ro: RM ANOVA *F*_(1.8, 12.6)_=0.1, *P*=0.9). *P* values indicate Holm-Sidak post-hoc comparisons. (D) Over the drug administration period, contralateral (*t*_(14)_=3.2) and ipsilateral (*t*_(14)_=2.5) cortical responses changed significantly less in Ro 25-6981-than vehicle-treated animals (2-tailed *t* tests). There was no effect on the ODI (Mann-Whitney test, U=27). Data are shown as mean±SEM. N indicates number of mice.

**Figure 3.**
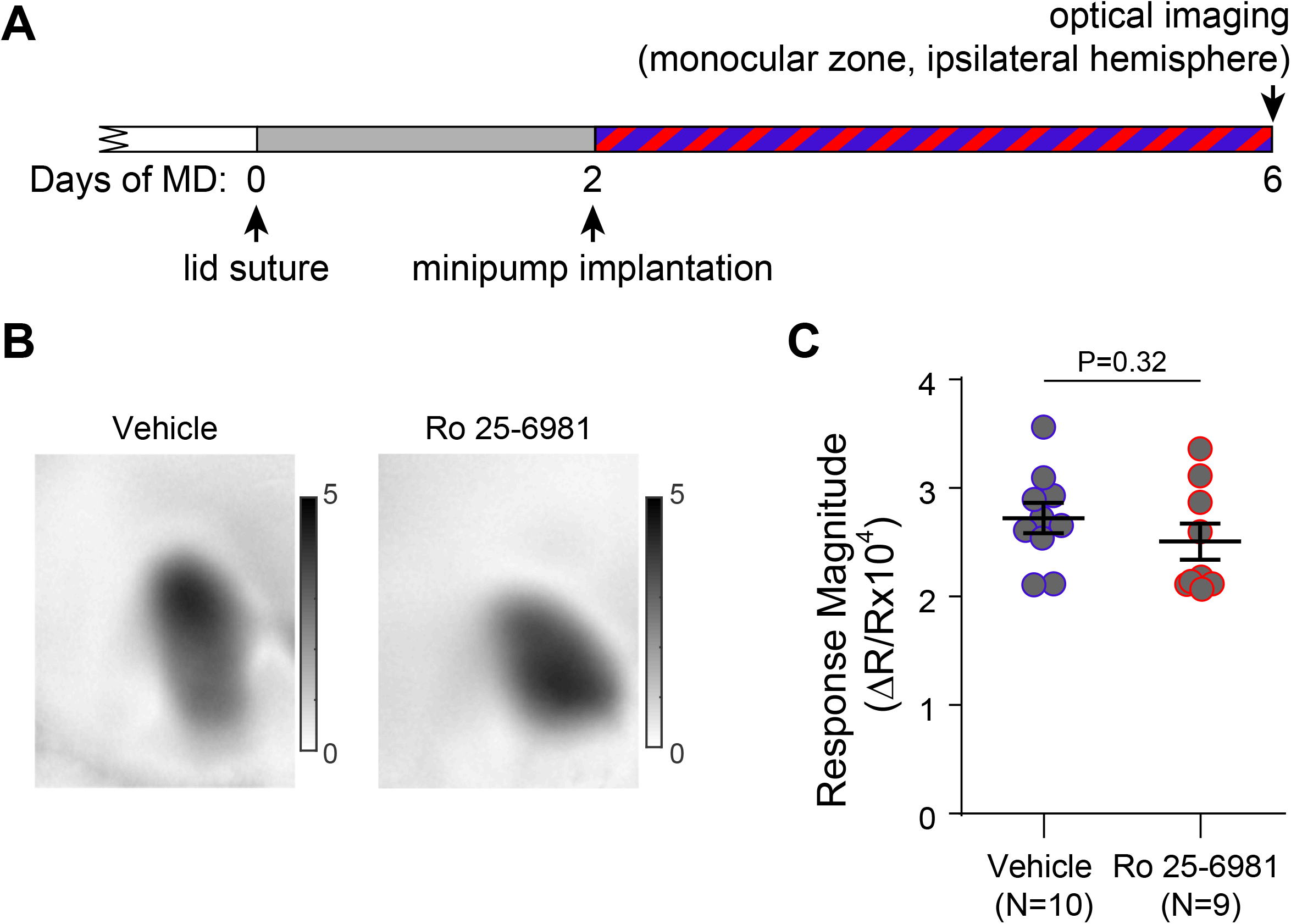
Ro 25-6981 does not affect map strength in non-remodeling V1. (A) Experimental design. Monocular cortex in the control hemisphere (ipsilateral to the deprived eye) was imaged while visually stimulating the monocular zone of the open eye. (B) Representative maps. (C) 4d of Ro 25-6981 treatment did not decrease map strength (*t* test *t*_(17)_=1.01). Data are shown as mean±SEM and N indicates number of mice.

### GluN2B is required for depression during early monocular deprivation

6d MD increases the GluN2B component of NMDA receptor current in V1 (Guo et al., 2017). We tested whether shorter (2d) MD was also sufficient to change NMDA receptor composition at visual cortical synapses. We recorded evoked NMDA receptor currents in the V1 layer 4-2/3 pathway and found no difference in the decay constant between deprived and non-deprived cortices in either V1m or V1b (Fig. 4A, B), indicating that brief MD does not increase the GluN2B component of the NMDA receptor current.

**Figure 4.**
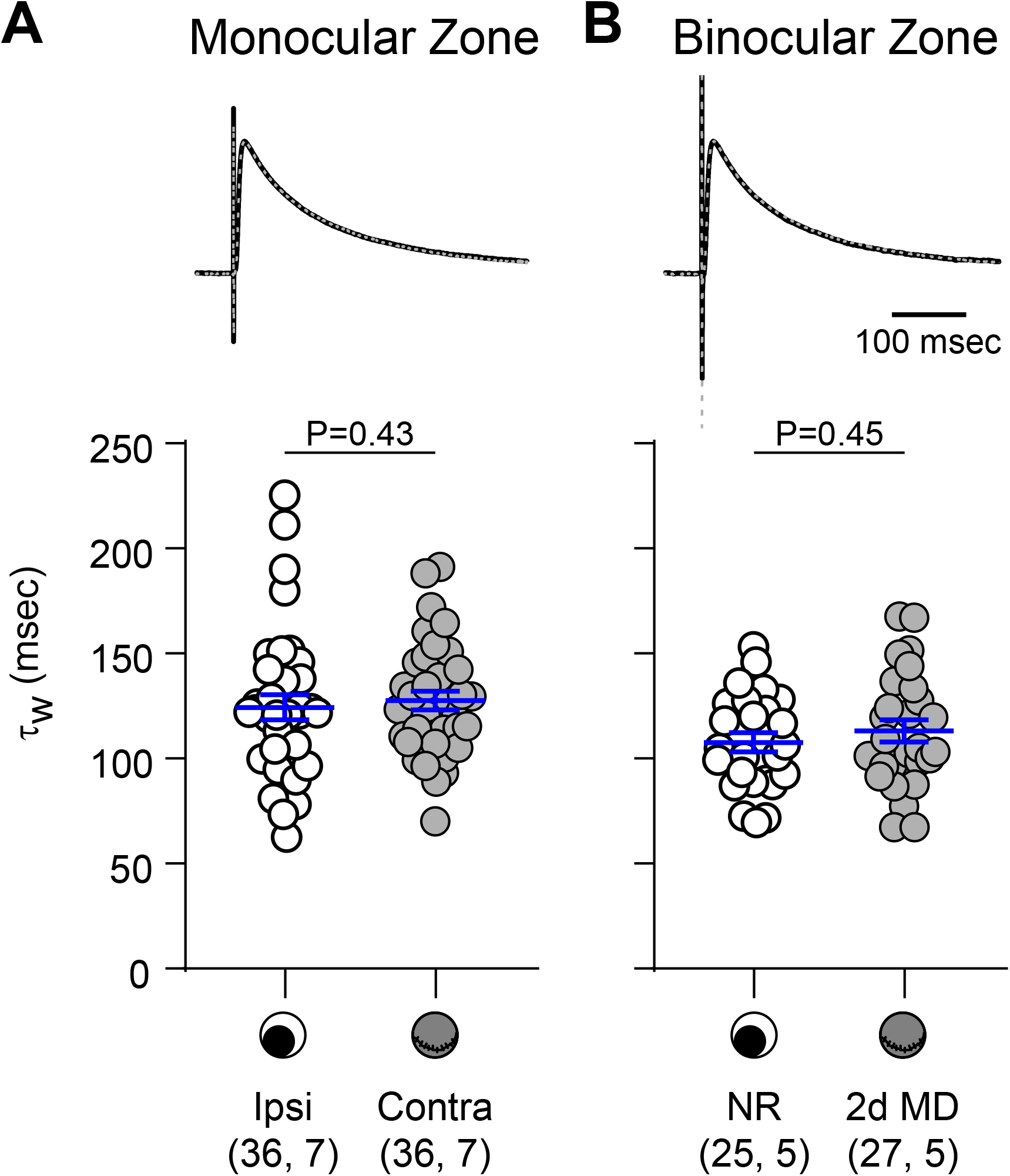
The GluN2B component of the NMDA response does not increase over 2d MD. (A) Top: Average evoked NMDA receptor responses in the monocular zone of control (ipsilateral; black) and deprived (contralateral; dashed grey) hemispheres obtained from the same animals. Bottom: The decay constant did not differ between hemispheres (Mann-Whitney *U*=577). (B) Top: Averaged evoked NMDA receptor responses in the binocular zone of normally reared (NR) animals (black), and in the deprived hemisphere of a second group of animals after 2d MD (dashed grey). Bottom: The decay constant did not differ between conditions (*t* test *t*_(50)_=0.8). Data are shown as mean±SEM and sample size is indicated as (cells, mice).

Although NMDA receptor subunit composition did not change with 2d MD, pre-existing GluN2B-containing NMDA receptors at the synapse may be required for decreased synaptic strength and visual response amplitude following brief MD. We therefore tested the effects of GluN2B-specific blockade during brief MD in both V1m and V1b (Fig. 5A). We first recorded mEPSCs in V1m of acute slices from mice that received continuous vehicle or Ro 25-6981 administration during 2d MD. In the vehicle group, mEPSC amplitude was significantly smaller in the deprived than in the nondeprived hemisphere (Table 2). In contrast, GluN2B blockade prevented mEPSC amplitude from decreasing in the deprived hemisphere (Fig. 5B, C; Table 2). Second, we measured the amplitude of visually evoked responses in V1b using optical imaging of the intrinsic cortical signal. Consistent with V1m, blocking GluN2B-containing NMDA receptors prevented the decrease in deprived eye response magnitude and the resulting decrease in ocular dominance index (ODI) (Fig. 5 D, E).

**Table 2.**
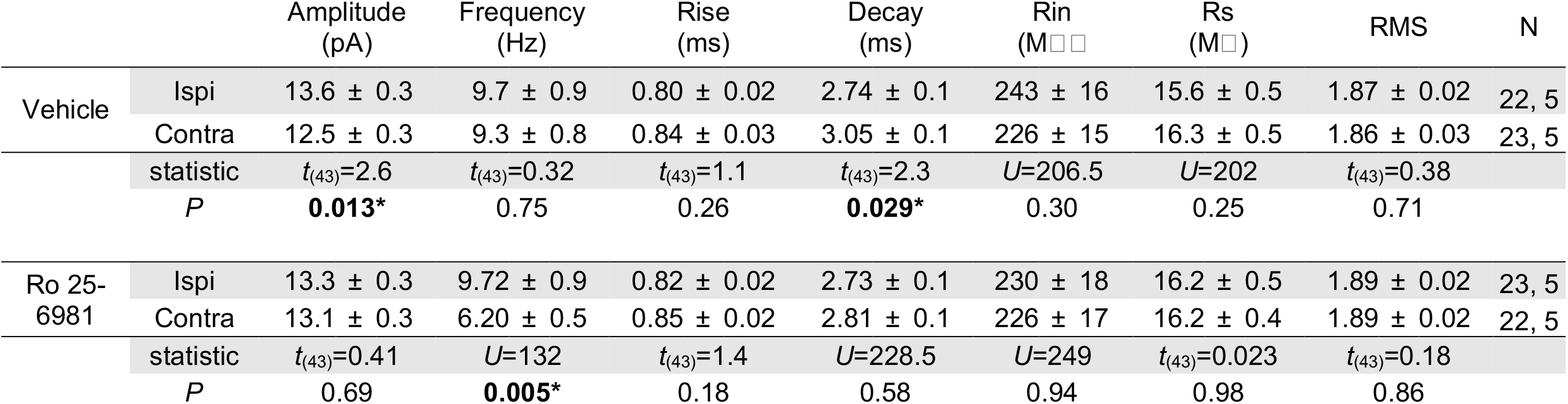
mEPSC properties and recording conditions after 2d MD (related to Figure 3). For each treatment, data were compared between control (ipsi) and deprived (contra) hemispheres using 2-tailed unpaired *t* tests (*t* statistic reported) or Mann-Whitney tests (*U* statistic reported), as indicated. Bold values indicate significant differences. Data are shown as mean±SEM. N is reported as cells, mice.

**Figure 5.**
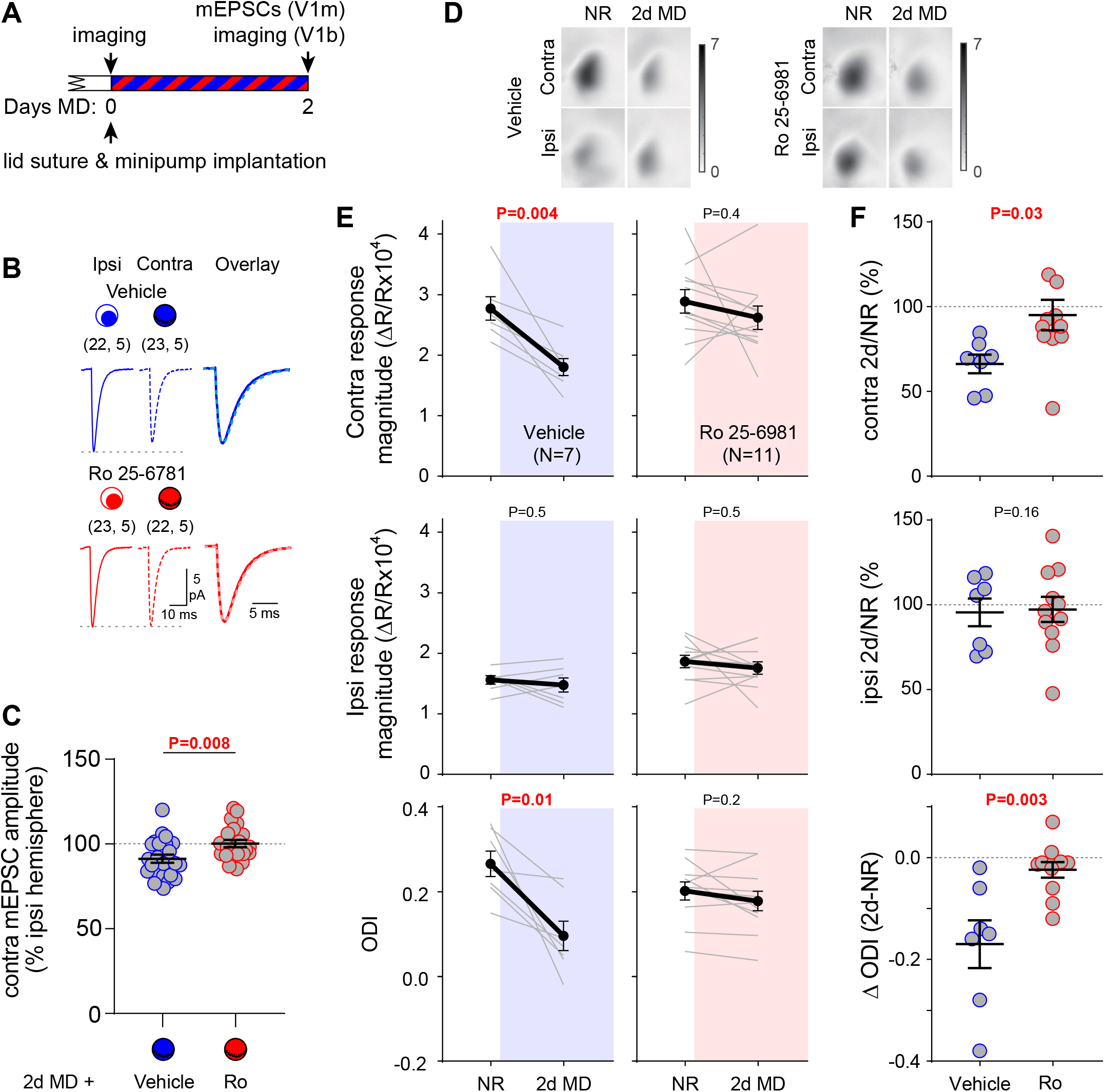
GluN2B-containing NMDA receptors are required for depression of deprived eye responses following brief MD. (A) Experimental design. Mice were implanted subcutaneously with a minipump containing Ro 25-6981 or its vehicle and the right eyelid was sutured shut for two days. (B) Averaged mEPSC traces, recorded in the monocular zone of each hemisphere. Sample size is indicated as (cells, mice). (C) mEPSC amplitude in the contralateral hemisphere is expressed as percentage of ipsilateral hemisphere in the same animal. GluN2B blockade prevented the decrease in mEPSC amplitude (*t* test *t*_(43)_=2.8). Data are shown as mean±SEM. Kinetics and recording conditions are reported in Table 2. (D) Representative images of visual responses in the binocular zone, measured longitudinally by optical imaging of the intrinsic cortical signal. (E) Contralateral eye response magnitude and ODI decreased in vehicle (Contra: *t*_(6)_=4.5; ODI: *t*_(6)_=3.6) but not Ro 25-6981 (*t*_(10)_=1.0, *t*_(10)_=1.5) treated mice over 2d MD. Ipsilateral eye responses were unchanged in both groups (Vehicle: *t*_(6)_=0.7; Ro *t*_(10)_=0.7). Comparisons were made using 2-tailed paired *t* tests. (F) The decrease in contralateral eye response and ODI over the drug administration period was greater in vehicle-than Ro 25-6981-treated animals (Contra: *t*_(16)_=2.4; ODI: *t*_(16)_=3.5). Change in ipsilateral eye response did not differ between groups (*t*_(16)_=0.16). Comparisons were made using 2-tailed *t* tests. Data are shown as mean±SEM and N indicates number of mice.

## Discussion

Here, we show that GluN2B-containing NMDA receptor activity is required for both phases of ODP: the initial depression of deprived eye responses and delayed potentiation of non-deprived eye responses. During the initial phase, levels of synaptic GluN2B-containing NMDA receptors are unaltered, and pre-existing GluN2B is required for synaptic weakening. Longer MD increases synaptic GluN2B levels, which favors Hebbian LTP (Guo et al., 2017; Philpot et al., 2001). These GluN2B-containing receptors are required for delayed synaptic strengthening, indicating that decreased LTP threshold, rather than NMDA receptor-independent synaptic scaling, underlies homeostasis in this model.

These results are consistent with the interpretation that NMDA receptor-dependent Hebbian LTD drives the initial synaptic weakening in V1 during ODP (Fong et al., 2020; Heynen et al., 2003; Lambo and Turrigiano, 2013). Our results further demonstrate a role specifically for GluN2B-containing NMDA receptors in the weakening of deprived eye responses, in line with LTD mechanisms in hippocampus and timing-dependent LTD in V1 (Fox et al., 2006; Ge et al., 2010; Guo et al., 2012; Izumi et al., 2006; Liu et al., 2004). Although GluN2B-containing NMDA receptors are necessary for LTD during ODP, additional mechanisms, such as endocannabinoid signaling, are also required (Liu et al., 2008); how these mechanisms interact to weaken deprived eye responses remains to be seen.

Homeostatic strengthening via the sliding threshold occurs during multiple types of visual deprivation, including MD (Fig. 1, 2) and dark exposure (Bridi et al., 2018). During dark exposure, the sliding threshold is engaged by altered firing patterns, whereas synaptic scaling is thought to be caused by changes in overall spiking and/or synaptic activity (Bridi et al., 2018; Fong et al., 2015; Turrigiano et al., 1998). Like dark exposure, MD has moderate effects on overall firing rates in V1, supporting the idea that homeostasis during visual deprivation depends more on activity patterns than spike rate (Aton et al., 2013; Fiser et al., 2004; Hengen et al., 2013; Torrado Pacheco et al., 2019). In contrast, synaptic scaling depends on dramatic reductions in firing rate that can be achieved by pharmacological or chemogenetic interventions *in vivo*, in line with the hypothesis that distinct homeostatic mechanisms operate within different activity ranges (Bridi et al., 2018; Wen and Turrigiano, 2021).

### Mechanisms of homeostasis

Strengthening of visual responses during sensory deprivation requires molecular mechanisms directly involved in metaplasticity and LTP, including GluA1-containing AMPA receptors and (GluN2B-containing) NMDA receptors (Fig 2; Ranson et al., 2013; Rodriguez et al., 2019; but see Toyoizumi et al., 2014). However, several mechanisms implicated in synaptic upscaling are also required, warranting closer examination of how their complex roles *in vivo* might contribute to homeostasis.

Multiple types of visual deprivation cause rapid cortical disinhibition onto principal cells (Aton et al., 2013; Gao et al., 2017; Hengen et al., 2013; Kuhlman et al.; Severin et al., 2021; van Versendaal et al., 2012). The altered activity patterns that result from disinhibition (due to increased spontaneous, decorrelated firing) are thought to slide the plasticity threshold in favor of LTP (Bridi et al., 2018). Retinoic acid and tumor necrosis factor α signaling can both mediate disinhibition (Pribiag and Stellwagen, 2013; Zhong et al., 2018). Therefore, these mechanisms may contribute to homeostasis during sensory deprivation *in vivo* by promoting LTP, distinct from their roles in scaling *in vitro* (Aoto et al., 2008; Kaneko et al., 2008; Stellwagen and Malenka, 2006). It would be of great interest to determine whether these mechanisms are required for the increase in GluN2B levels during sensory deprivation.

In light of our findings we speculate that, once the plasticity threshold is lowered, LTP maintenance is required for the continued expression of homeostasis during sensory deprivation. Some molecular processes play dual roles in LTP maintenance and scaling, which may explain why mechanisms required for scaling *in vitro* are also necessary for homeostasis during sensory deprivation *in vivo*. For example, interfering with GluA2-mediated AMPA receptor trafficking and activity regulated cytoskeleton-associated protein (Arc) prevent homeostasis during MD (Lambo and Turrigiano, 2013; McCurry et al., 2010). These findings have been attributed to the roles of GluA2 and Arc in synaptic scaling (Gainey et al., 2009; Gao et al., 2010; Shepherd et al., 2006), but may instead be due to their involvement in LTP maintenance (Guzowski et al., 2000; Plath et al., 2006; Shi et al., 2001). In line with this hypothesis, Arc KO mice are also deficient in stimulus-selective response potentiation, a form of visual cortical plasticity that engages LTP mechanisms (Frenkel et al., 2006; McCurry et al., 2010).

The current study in juvenile mice reveals multiple roles for GluN2B-containing NMDA receptors in visual cortical plasticity. Molecular mechanisms of synaptic weakening and homeostatic strengthening in V1 vary across developmental stages (Ranson et al., 2012, 2013), cortical layers (Crozier et al., 2007; Fong et al., 2020; Liu et al., 2008), and neuronal subpopulations (Pandey et al., 2022). Determining how GluN2B interacts with each of these distinct processes will shed light on how the cortex adapts to altered sensory input across the lifespan.

